# Genome‑wide identification and characterization of Transcription Factors of Basic Leucine Zipper Family in *Malus domestica*

**DOI:** 10.1101/075994

**Authors:** Zhengrong Zhang, Haoran Cui, Shanshan Xing, Xin Liu, Xuesen Chen, Xiaoyun Wang

## Abstract

Basic leucine zipper proteins (bZIP) contain a basic DNA-binding region and a leucine zipper region, acting as transcriptional factors in regulation of gene expression exclusively in eukaryotes. In this investigation, total 116 bZIP members were identified in apple genome and mapped on all 17 chromosomes with various densities as *M*.bZIPs. All these members were divided into six groups according to the phylogenetic relationship combining with bZIPs from rice and *Arabidopsis*. Investigating gene structure of *M*.bZIPs, five splicing patterns of intron were found in the DNA-binging region with no splicing position and splicing positions at different nucleotide of codons or different positions. Analyzing of protein structure of *M*.bZIPs, twenty-five motifs were identified with certain characteristic in different phylogenetic groups. To predict dimerization of leucine zipper region, the key positions of amino acid in heptad(s) were investigated. The results showed that most *M.*bZIPs may form hetero-dimer or homo-dimer and some *M*.bZIPs may form both. Expression experiment revealed that *M*.bZIP genes have organ-specific expression and widely expressed in flowers, leaves, and fruits. To investigate the response of *M.*bZIPs to abiotic stresses, the promoter sequences of randomly selected *M*.bZIP genes were analyzed. *Cis*-acting elements related to multiple stresses were found existing widely in promoter sequences. Quantitative real-time PCR results further demonstrated that the expression of some *M*.bZIP genes were quite sensitive to exogenous abscisic acid and osmotic treatments.

## 1. Introduction

As a large family, basic leucine zipper proteins (bZIP) are widely present in eukaryotes (Hurst 1994). There are different bZIP members in different species, for example, 17 in *Saccharomyces cerevisiae*, 31 *in Caenorhabditis elegans*, 27 in *Drosophila* (Fassler et al. 2002), 56 in human (Vinson et al. 2002). Generally, bZIP members in plants are much more, 75 in *Arabidopsis* (Jakoby et al. 2002), 89 in rice (Nijhawan et al. 2008), 92 in *sorghum* (Wang et al. 2011) and 131 in soybean (Liao et al. 2008). More bZIP members in plants may suggest that bZIPs are more important in plants than in animals. As transcription factor, bZIP binds to negatively charged DNA in DNA-binding region while the leucine zipper region formed homo- and/or heter-dimer to maintain the stability of the combination. The DNA-binding region is highly conserved, containing about 18 amino acid residues (Asn-X_7_-Arg/Lys-X_9_). Leucine zipper region is made up of hepated(s) with repeated leucine or similar hydrophobic amino acids. The amino acids at positions nearing interface of leucine zipper determine the dimerization stability and specificity (Vinson et al. 2002, Newman and Keating 2003).

Plant bZIPs is reported to be involved in diverse physiological processes, especially responses to abiotic/biotic stresses. In maize, *Zm*bZIP17 functions as a TF by interacting with ABA-responsive *cis*-elements (ABRE) of promoter in gene, acting as an stress transducer in endoplasmic reticulum (Yang et al. 2013). *Os*bZIP71 is also strongly induced in ABA-mediated drought and salt tolerance in rice (Liu et al. 2014). Compared to animal, various mechanism of anti-stress is more important, since plants are non-mobility. Although some plant bZIPs were verified and some investigations on their function have been reported. However, such studies have mostly focused on herbaceous plant species, and the bZIP genes of woody economically important fruit tree species have received less attention

Apple (*Malus domestica*) is one of the most economically important woody plant and fruit crops in the world. Recently, the divergence of the bZIP gene family in apple, strawberry and peach were reported. It showed that bZIP genes exist multiple modes of gene evolution after duplication (Wang et al. 2015). Using iTAK-Plant Transcription factor and Protein Kinase Identifier and Classifier database, Zhao et al. obtained 112 apple bZIP members and divided all these members into 11 groups. Detailed investigation of some members in one group (group A) indicated that they had organ-specific expression and were responsive to high salinity and drought, as well as to different phytohormones (Zhao et al. 2016).

Studies on bZIP family from different species indicated that gene structure and splicing patterns of intron are important for their functional evolution, since it would affect the coding sequences of protein, especially for the functional basic domain (Liu and Chu 2015). As TFs, bZIP regulate the expression of other genes by binding DNA, which functional motifs are quite important to their function. Dimerization is also an important functinal characteristic of bZIP protein which determines the stability of the binding between bZIP transcription factor and DNA.

In the present study, total 116 *M.*bZIP encoding genes were identified from the apple genome and divided them into different groups based on phylogenetic relationship. Splicing patterns of intron in DNA-binding region and characterized dimerization property within the leucine zipper regions were analyzed. As protein structure, different functional motifs were identified, some motifs were involved in ABA-response signal-transduction pathway and abiotic stress response. C*is*-acting elements related to abiotic stress tolerance and hormone signaling were massive existence in the promoters of *M.*bZIP genes. The expressions of some *M.*bZIP genes which were randomly selected in different phylogenetic groups were quite responsive to the ABA and osmotic treatments.

## 2. Results

### 2.1. Identification, Chromosomal Location and Expansion Patterns of bZIP Genes in Apple

To identify bZIP proteins in apple genome, two methods were used. For the first method, all of the identified sequences of bZIP proteins from *Arabidopsis* and rice were used as queries to search the Golden Delicious peptide database (GDR http://www.rosaceae.org/) using software BLASTp (Gagne et al. 2002). Trying not to miss potential proteins and taking account of credible results, the E value cutoff was set to 0.001 as in references of similar investigations (Li et al. 2011). From this method, about 700 proteins were screened out. For the second method, Hidden Markov Model (HMM) profile of the bZIP domain from characteristics of the amino acid sequences established by the pfam website were used as a query to search in the Golden Delicious peptide database. Total 199 proteins were obtained from this method. To further identify whether all above proteins contains the bZIP domain, the sequences were submitted to the Interpro Database to be verified. Finally, 116 proteins have been demonstrated to have bZIP domain. The detail information of these proteins with their relative encoding genes was listed in Table 1.

**Table 1.**
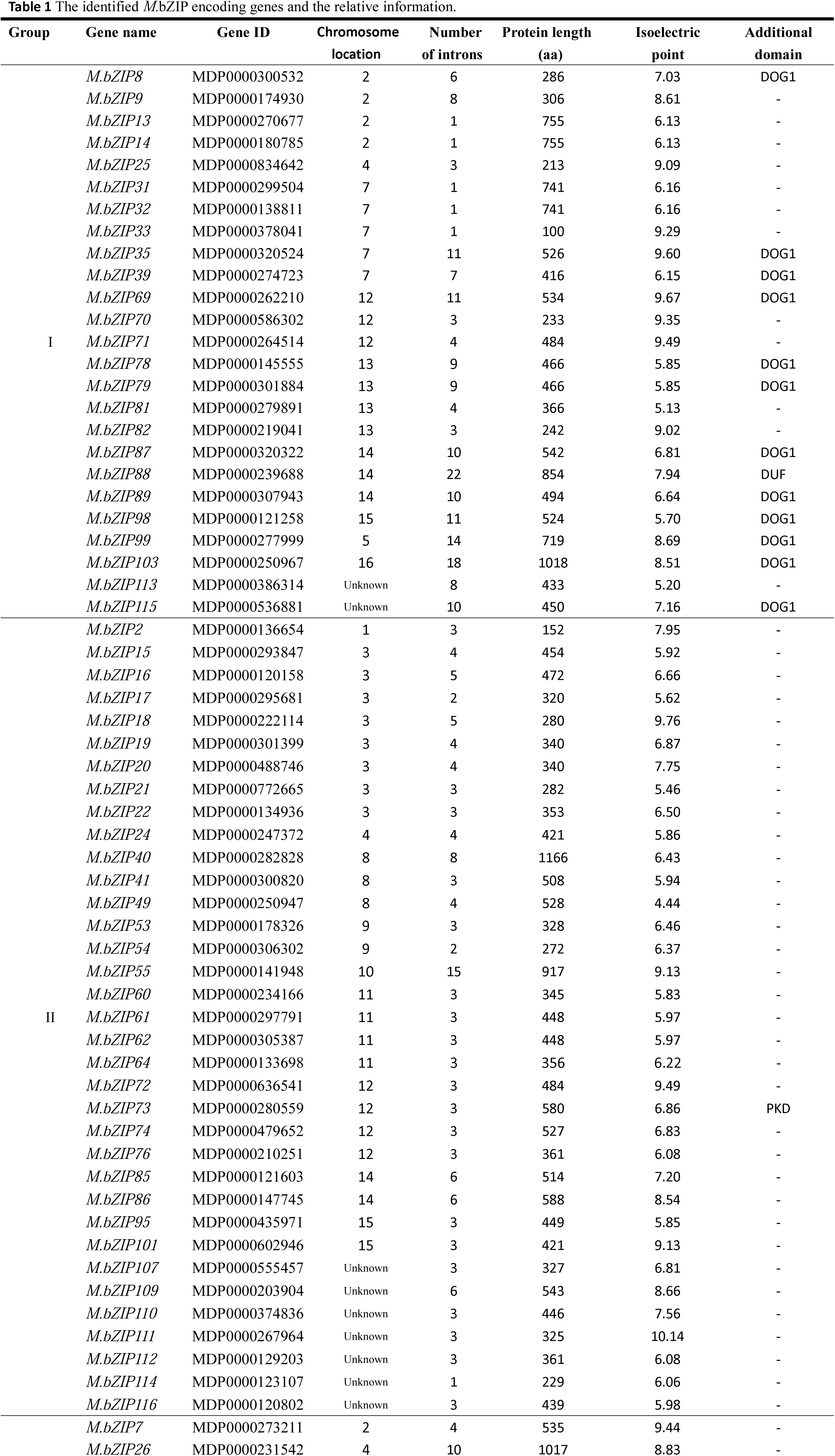
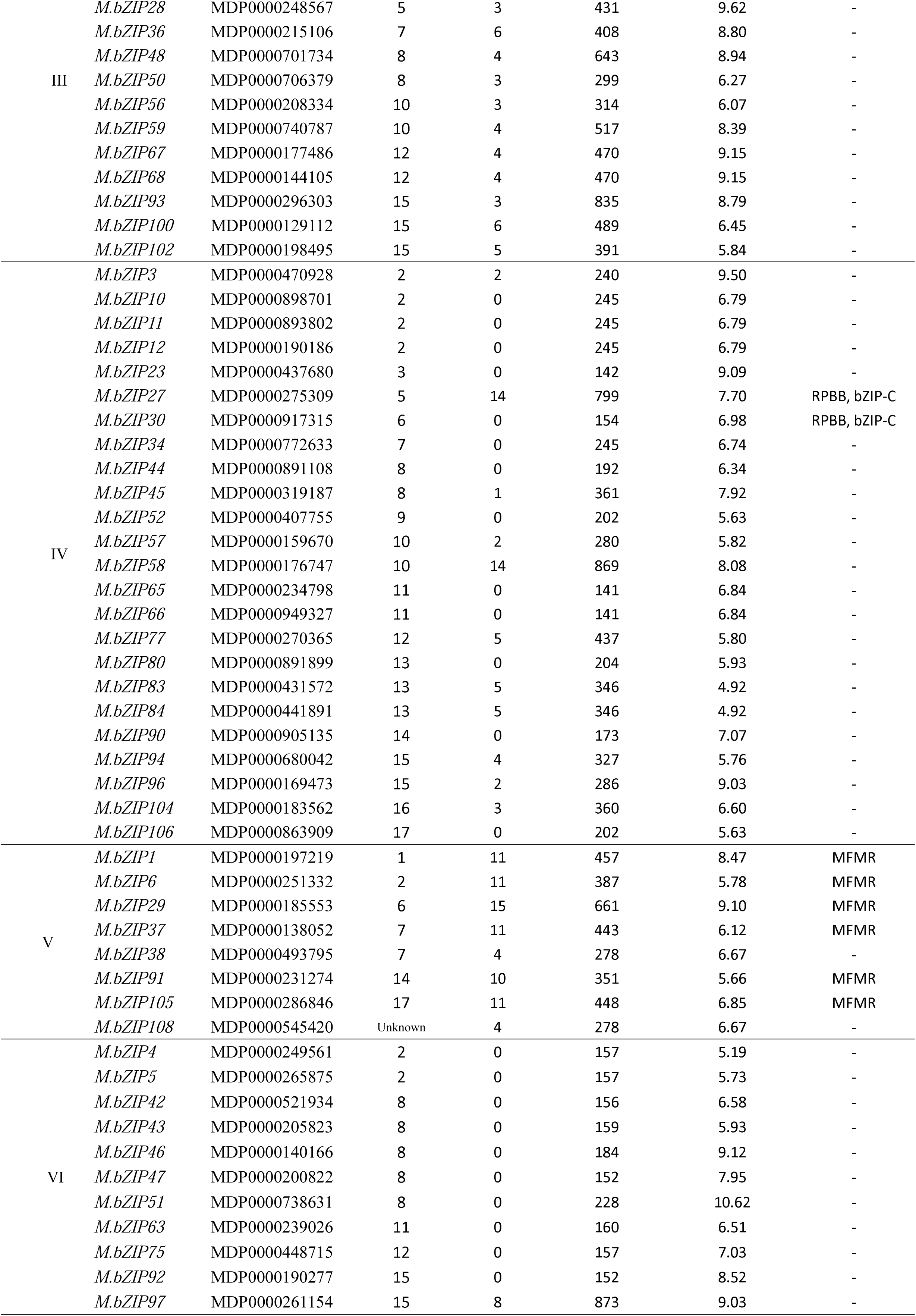
The identified *M.*bZIP encoding genes and the relative information.

For mapping 116 *M*.bZIP genes on the apple chromosomes, their gene ID and location data were retrieved from genome annotations downloaded from the Genome Database for Rosaceae (GDR http://www.rosaceae.org/). The chromosome map showing the physical location of all encoding *M.*bZIP genes (for short *M.*bZIP genes) was generated using procedure of MapDraw (Figure 1). Most *M.*bZIP (107 of 116) were mapped on all 17 apple chromosomes and designated as *M.*bZIP 1-107 according to their top-to-bottom position on chromosomes from 1 to 17, while nine genes could not be mapped on any chromosome and designated as *M.*bZIP108-116. Presumably these nine genes could be localized to unassembled genomic sequence scaffolds (Li et al. 2011). The number of *M.*bZIP genes on different chromosomes was variable. There were four chromosomes harboring low density of *M.*bZIP genes, chromosome 1, 5, 6 and 17 with only two *M.*bZIP genes on each chromosome. Other chromosomes had relatively high density and *M.*bZIP genes usually emerged with gene clusters on each chromosome. Further analyzed the distance between genes which located in the cluster, within 30kb of each other was defined as tandem duplication gene (Du et al. 2013). These tandem duplicates genes were highlighted with green block as shown in Figure 1 and listed in Table S1. Based on this result, we hypothesized that the tandem duplication of genes was one reason for the expansion of the *M.*bZIP genes. In addition to gene tandem duplication, a genome-wide duplication (GWD) event also occurred in apple and formed several intra-genomic homologous regions (Li et al. 2011). Large numbers of *M.*bZIP genes were located in homologous regions (gray shadow region in Figure 1) on different chromosomes, which suggest that genome duplication was another important way for expansion of *M.*bZIP genes.

**Figure 1.**
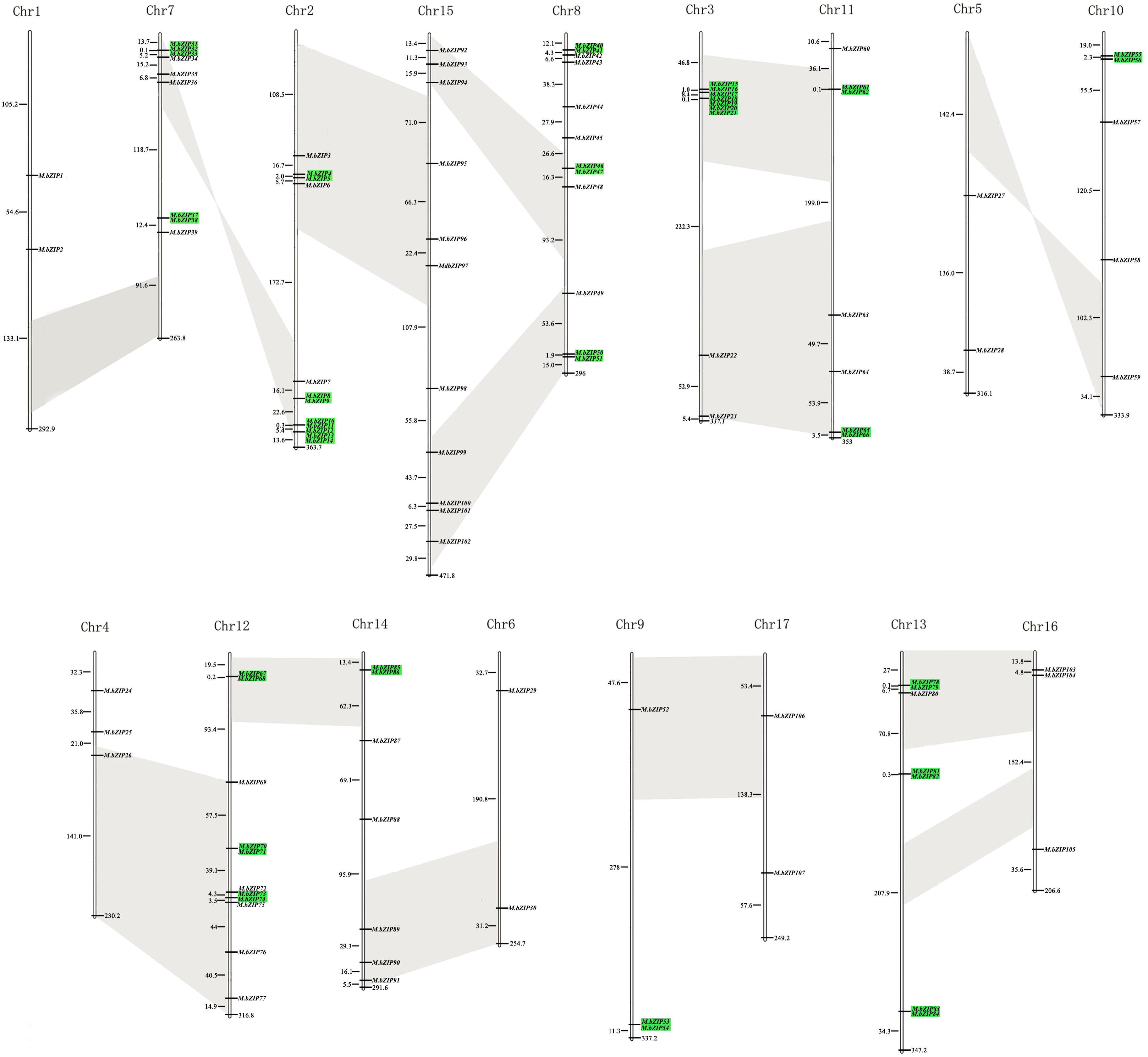
Distribution of the *M.*bZIP genes in 17 apple chromosomes. The numbers to the left of each chromosome represent a megabase between two *M.*bZIP genes. Tandem duplications are indicated by light green, gray block represent intragenomic homologous regions.

### 2.2. Phylogenetic Analysis and Classfication of the *M.*bZIP family

Previous studies have shown that the evolution of *bZIP* genes was conservative in plants (Liu and Chu 2015). To obtain the credible phylogenetic relationship of the *M.*bZIP proteins, bZIP members from apple (116), *Arabidopsis* (72) and rice (89) were used together to construct a phylogenetic tree (Figure 2). Comparing evolutionary grouping of *Arabidopsis* and rice from previous reports, the phylogenetic tree constructed in this investigation was credible. All *M.*bZIP members can be divided into six groups, designated as group I to VI, with 25, 35, 13, 24, 8 and 11 members in each group, respectively. The group II was the largest group with 35 members, and the group V was the least with only 8 members. Most of the *M.*bZIP genes exhibited closer relationship with *At*bZIPs than *Os*bZIPs and they were hierarchical clustered in the same clade in the phylogenetic tree, which suggested that dicots of apple and *Arabidopsis* might had the similar patterns of their bZIP families in evolutionary.

**Figure 2.**
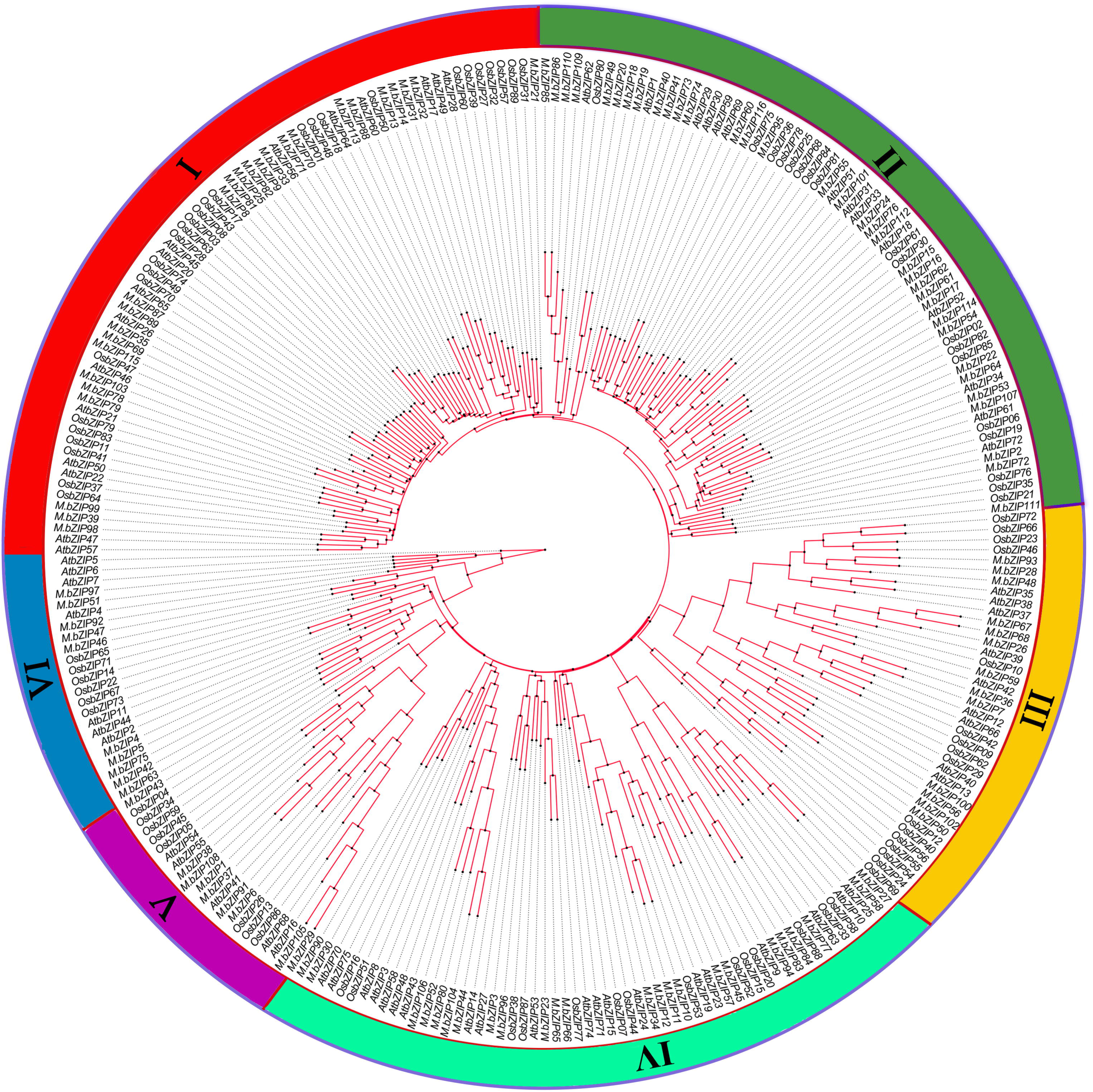
The phylogenetic tree of bZIP proteins from apple, rice and *Arabidopsis*. It was constructed by aligning sequences of bZIP protein from apple (116) *Arabidopsis* (72) and rice (89).

### 2.3. Gene Structure of *M.*bZIP genes

As a result of key events in evolution, gene structure including the number and distribution of exon/intron, offers insights into understanding the emergence and evolution of a given gene (Patthy 1987). The exon/intron organization of 116 *M.*bZIP was analyzed based on the number, distribution and splicing patterns of intron. In total, 23 of 116 *M.*bZIP genes contained no intron and 93 of 116 *M.*bZIP genes contained varied introns, the number from one to twenty-two (Table 1). The members of *M.*bZIP genes in each group possessed similar gene structure in intons number and distribution. The number of introns in each group was uneven, with intronless genes only found in group IV and VI, dramatically varied from one to twenty-two introns in the group I. In the group II, most genes had less than six introns, and in the group V, most genes had more than ten introns (Table 1).

Further analyzing splicing patterns of DNA sequence encoding the conservative DNA-binding region (basic and hinge region) of bZIPs, five splicing patterns of intron were characterized and designated as pattern S-I to S-V (Figure 3 and Table S2). The pattern S-I was the most prevalent and lacked intron in this regions with 43 *M.*bZIPs from group IV and VI. The pattern S-II had one intron in phase 2 (splicing occurred between the second and the third nucleotide in one codon) in the basic region, while the pattern S-III had one intron in phase 0 (splicing occurred between codons) located between amino acid Gln and Ala. The members in pattern S-III was completely corresponded to that from group III and V except *M.*bZIP100. Pattern S-IV had two introns (each in phase 0), one located between amino acid Lys and Arg within the basic region and the other located between amino acid Gln and Ala within the hinge region, which member was exclusive in group I. The pattern S-V had only one intron in phase 1 (splicing occurred between the first and the second nucleotide in one codon) in the hinge region. Gene structure showed that each group of *M.*bZIPs mostly shared the same intron/exon structural pattern and remained conserved during the course of evolution of *M*.bZIP family genes, which were very consistent with those in other plants.

**Figure 3.**
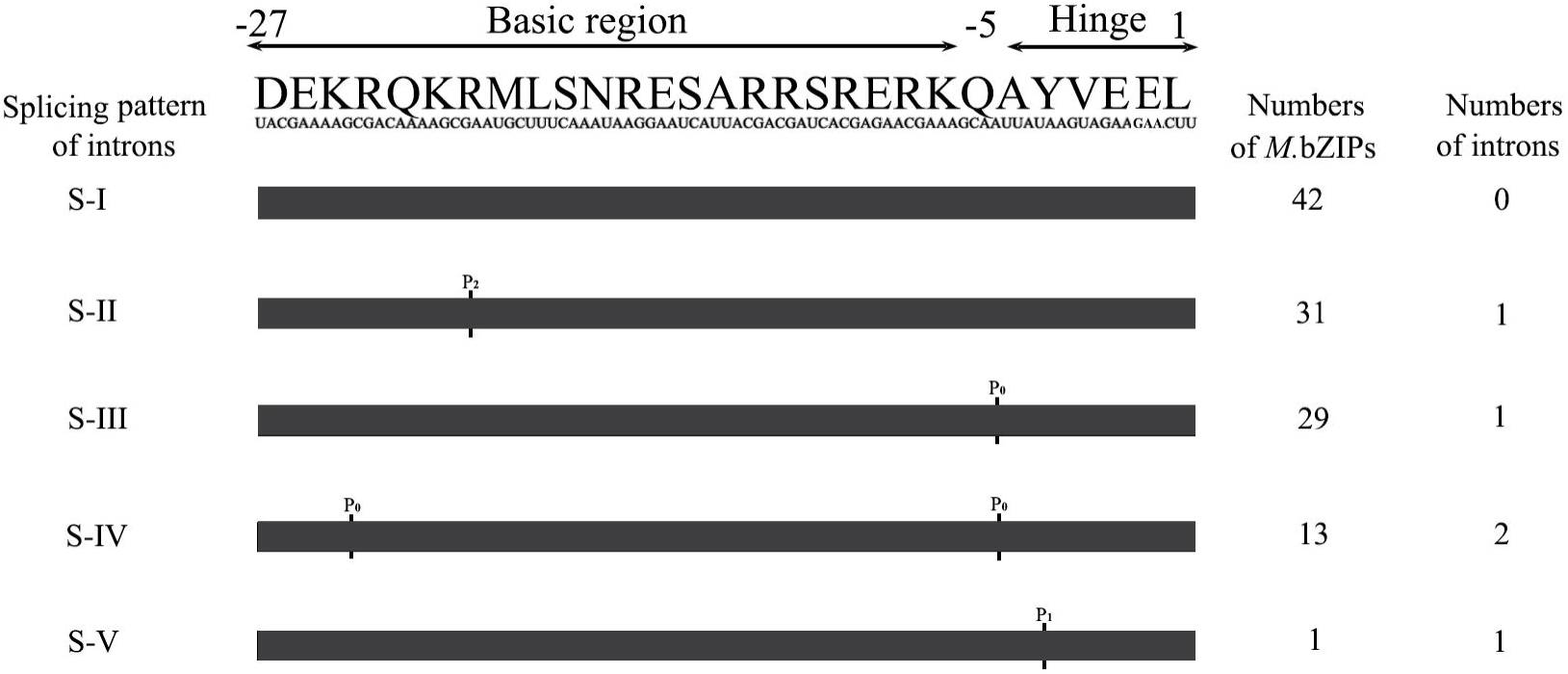
Splicing patterns of intron within the basic and hinge regions of the bZIP domains of the *M.*bZIPs. P_1_, P_2_ and P_0_ means splicing occurred after the first, second and third nucleotide of the condon, respectively.

### 2.4. Conserved Structural Features of *M.*bZIP Proteins

The bZIP domain is the core of the bZIP proteins, which preferentially binds to the promoter of their downstream target genes on specific *cis*-elements. To analyze the characteristics of the bZIP domain of *M.*bZIPs, the amino acid sequence logo of bZIP domain was generated by submitting all the sequences of 116 members to the MEME analysis tool (Figure 4). Most bZIP domain of *M.*bZIPs contained 39 amino acids and residues at position 11, 19, 29, 36 were much conserved as Asn, Arg /Lys, Leu and Leu. In all *M.*bZIPs, a conservative sequence could be identified from residue Asn (position 11) as Asn-X_7_-Arg/Lys-X_9_-Leu-X_6_-Leu (Table S3), which was accord with that of bZIP domains in other plants. Besides, some amino acids especially charged (Asp, Glu, Arg and Lys) were also conservative in this region (gray shadow in Figure 4).

**Figure 4.**
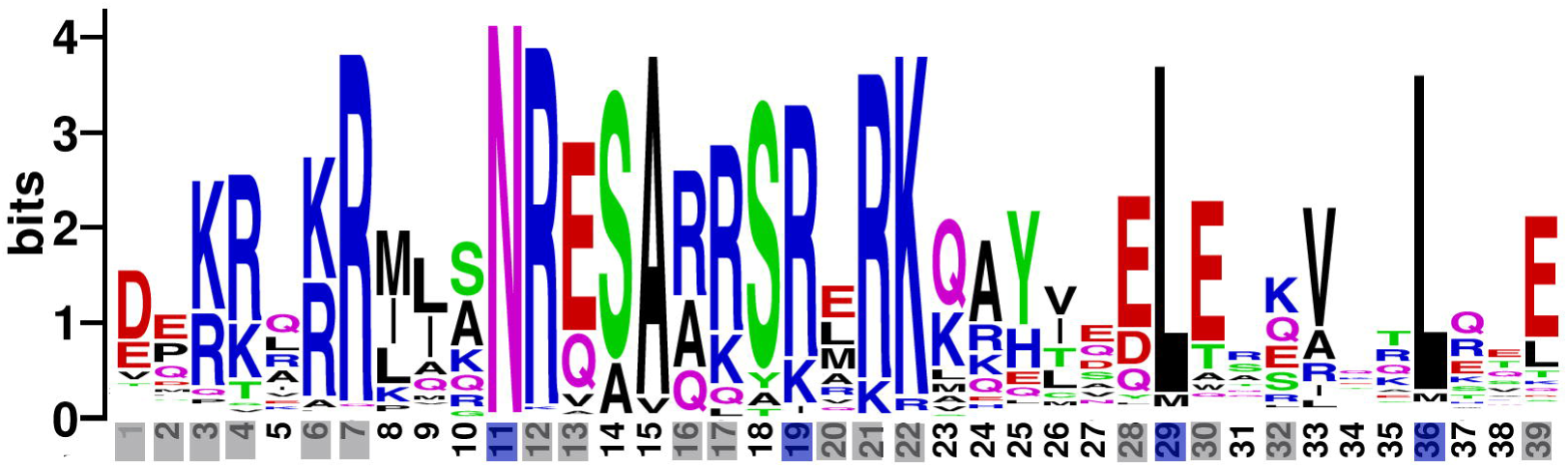
Sequence logos of bZIP domain from all *M.*bZIPs. Blue shadows represent the most conserved amino acids in the feature sequence of bZIP domain, gray shadows represent the charge amino acids of bZIP domain.

Besides the bZIP domain, bZIP proteins usually contain additional conserved motifs which might be related to the function of bZIP proteins as potential sites. A total of 25 motifs, including the conserved bZIP domain (be named motif 1) were identified in 116 *M*.bZIPs (Table S4) and their distribution was shown in Figure 5. Except motif 1, motif 7 was the most widely distributed in 69 *M.*bZIP members in each group. Some motif shared by several groups, motif 9 presented in four groups (group II, III, IV and V), motif 12 presented in three groups (group II, IV and VI), and motif 11 presented in two groups (group IV and V). Most of the conserved motifs, however, were presented in specific group, for example, motif 2, 3, 6, 14, 15 and 19 only existed in the group I, motif 4, 5, 13, 16, 20, 21, 23 and 24 only presented in the group II. Group-specific motifs may indicate their specific functions in each group.

**Figure 5.**
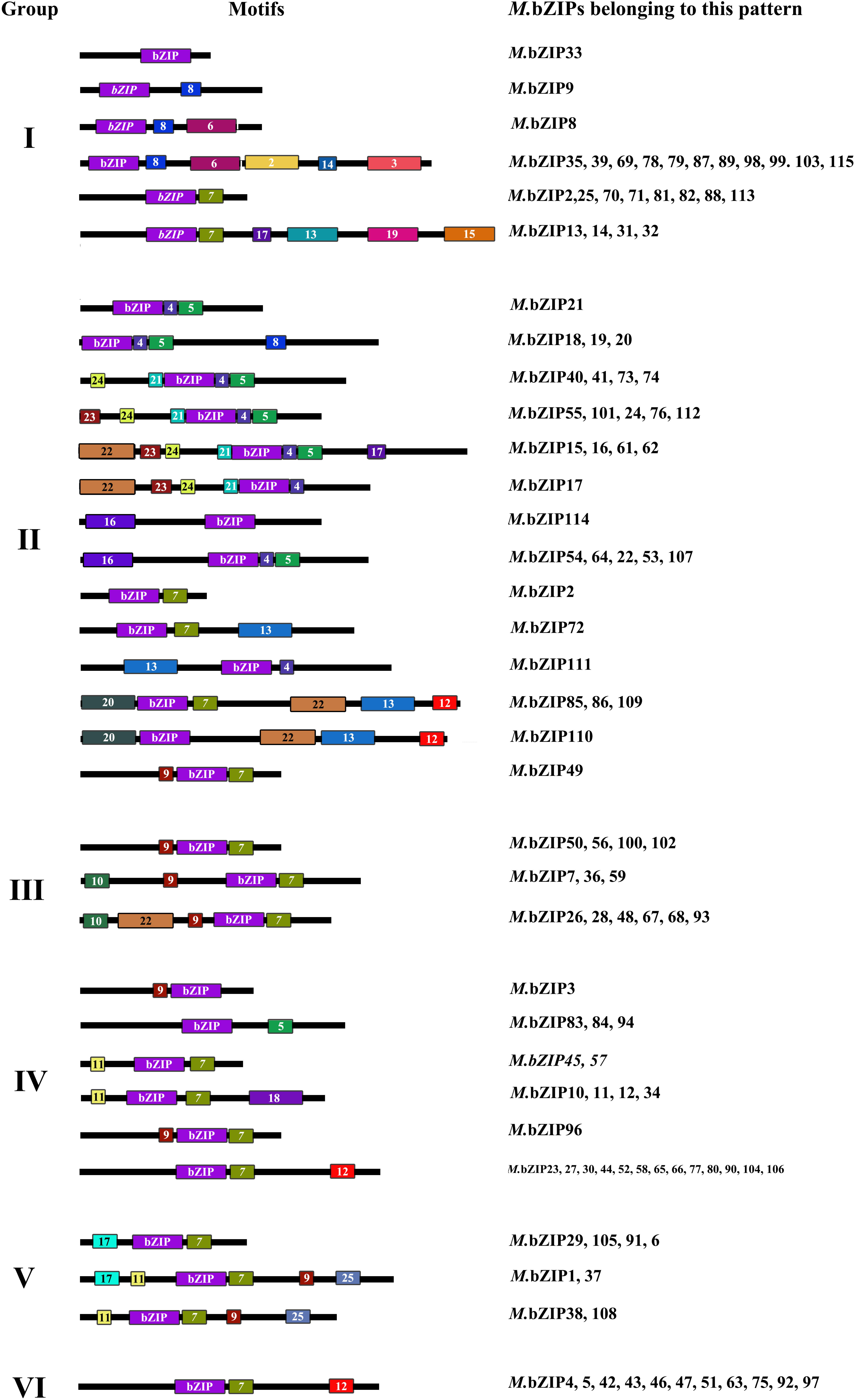
Protein structure analysis and motif identified in *M.*bZIPs. The bZIP domain is shown in purple block, other predicted motifs are indicated in different color blocks with numbers from 2 to 25. The details of predicted conserved motifs are given in table S3.

Protein phosphorylation, catalyzed by protein kinase is an important modification and directly related to protein function. Two conserved sequences, R/KxxS/T and S/TxxD/E, were verified as phosphorylated sites by a Ca^2+^ independent protein kinase and casein kinase II, respectively (Choi et al. 2000, Finkelstein and Lynch 2000, Nijhawan et al. 2008). Six motifs (motif 6, 8, 9, 12, 14 and 24) were verified harboring phosphorylation sites of casein kinase II. Nine motifs (motif 2, 3, 4, 11, 15, 16, 19, 20 and 23) harbored function site of Ca^2+^ independent protein kinase. Motif 3 and motif 18 contained both sites of above two kinases. Other functional sites were also identified in motifs. Motif 4 and motif 7 were extension of Leucine zipper region, function for the dimerization of *M.*bZIPs. Motif 6 only presented in the members of group I, was part of the DOG1 domain, which was related to the dormancy of the seeds (Bentsink et al. 2006). Motif 17 was found in six members of group V, with the presence of small stretches of Pro and aromatic amino acids Tyr, Phe and Trp, and is a part of the Pro-rich domain, similar motifs have been identified in arabidopsis, which have been shown to have transcription activation potential (Schindler et al. 1992).

### 2.5. Predicted Dimerization of *M.*bZIP proteins

bZIPs bind to DNA as dimer when acting as transcription factors (Deppmann et al. 2004). The leucine zipper region in bZIPs, arranged in the form of heptad repeats, is responsible for dimerization. Within each heptad, the amino acid positions are named as *a, b, c, d*, *e, f* and *g* in order in human and *Arabidopsis* (Vinson et al. 2002, Deppmann et al. 2004). Following this criteria, two to nine heptads were identified in different *M.*bZIPs (Table S5). Stability and specificity of leucine zipper dimmer are closely related to the amino acids present at the position *a, d, e* and *g*, since they are near the interface of leucine zipper (Wei et al. 2012). Amino acids at the position *a* and *d* are typically hydrophobic, forming a hydrophobic core is essential for interaction between two monomers. The position *e* and *g* usually present charged amino acids, which mediate the specificity of dimerization as well as stability (Nijhawan et al. 2008).

By analyzing all the 116 *M.*bZIPs, less than one tenth of charge amino acids (K, R, E and D) presented at position *a,* while other amino acids especially hydrophobic residues were predominant at the position *a* and *d* (Figure 6A). About 22% of amino acids present at the position *a* were Asn, equivalent amount with that of bZIP from maize (22%). Further analyzing the composition of each heptad, the highest frequency of Asn at the position *a* occurred both in the second and fifth heptads, accounting for 59% and 58% respectively (Figure 6B). The high frequency of Asn at the position *a* implied that it will form homo-dimerizing leucine zippers among the bZIP members, because Asn produce more stable N-N interactions than other amino acids (Nijhawan et al. 2008). Leucine present in position *d* was also import for dimerization stability. The frequency of Leu was high to 70% at the position *d* in *M.*bZIPs (Figure 6A), a similar level with that of bZIPs from rice (71%) and greater than that from *Arabidopsis* (56%).

**Figure 6.**
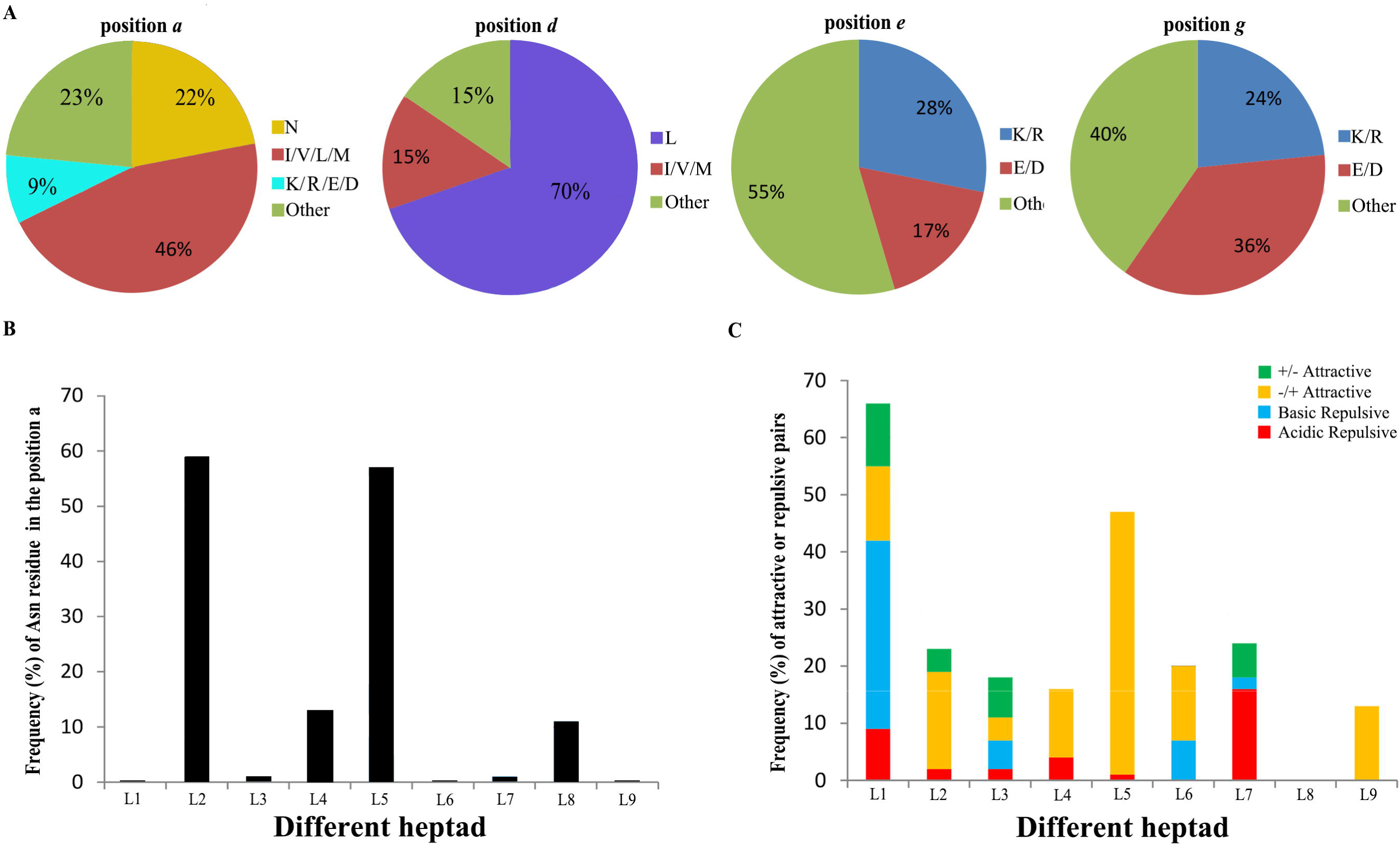
Prediction of dimerization properties of *M.*bZIPs. (A) The frequency of the amino acids at position *a, d, e* and *g*, these positions directly related to dimerization of the leucine zipper part of *M.*bZIPs. (B) The frequency of Asn residues present at the position *a* in the nine heptads for *M.*bZIPs. (C) The ratio of attractive or repulsive pair per heptad for *M.*bZIPs.

Different from the position *a* and *d*, the charged amino acids occupied near half in the position *e* and *g*, with frequencies of 45% and 60% respectively (Figure 6A).The attractive and repulsive *g* and *e* pairs governs dimerization properties of bZIPs (Deppmann et al. 2004). The amino acids present in each heptad of leucine zippers of *M.*bZIPs were characterized (Table S5) and the histogram of their frequency was presented in Figure 6C. Maximum frequency of the complete *g* and *e’* pairs appeared in the first heptad, showing basic repulsive in blue block. High frequency of attractive pairs were found in the second, fifth and ninth heptad, moreover, only -/+ attractive pairs were found in the ninth heptad.

Attractive pairs and presence of Asn in position *a* contribute to homo-dimerization, while repulsive and incomplete pairs, and charged residues in position *a* may favour hetero-dimerization (Wei et al. 2012, Liu and Chu 2015). According to these principles, total 116 *M.*bZIPs were classified into 29 sub-families (Table S5). One subfamily favoured homo-dimerization (subfamily BZ1 in Table S5), since attractive pairs presented in the first and second heptads and lacking any repulsive interactions. Two subfamilies could be considered having hetero-dimerizing specificity because they contained only repulsive interhelical interactions (subfamily BZ2 and BZ3). There was a subfamily (subfamily BZ29) was thought not to form dimer because these proteins lacked leucine zipper region. Other twenty-five subfamilies had both homo- and hetero-dimerization properties.

### 2.5. The Organ-Specific Expression of *M.*bZIP Genes

Evidences suggested that bZIPs are widely involved in the integration and development of many organs in plants, such as seed maturation and germination (Toh et al. 2012), floral induction and development (Walsh and Freeling 1999, Wigge et al. 2005). To further analyze the organ-specific expression of *M.*bZIP genes, GSE42873, an expression profile from the GEO DataSets were used. According to the expression of 116 *M.*bZIP genes in different organs, four expression types were divided and designated as type I-IV (Figure 7). Generally, most of *M.*bZIP genes were higher expression in flowers, fruits and leaves, indicating that *M.*bZIP genes widely participated in the process of flower development and fruit ripening. Type I contained sixteen members with low expression in seedlings. Compared with other types, the gene expression in this type was relatively high in the seeds indicating their potential function in seed germination. In type II, twenty-two genes were clustered together due to their highest expression in the flowers and lowest expression in the seeds. There were sixty-eight genes that were clustered into type III because they showed a low expression in the seeds, seedling, roots and stems compared with other organs. In type IV, all of the ten genes had a low expression in the seeds, but most of them had a high expression in the roots, stems and seedlings, implying that the gene of type IV involved in the vegetative growth of apple.

**Figure 7.**
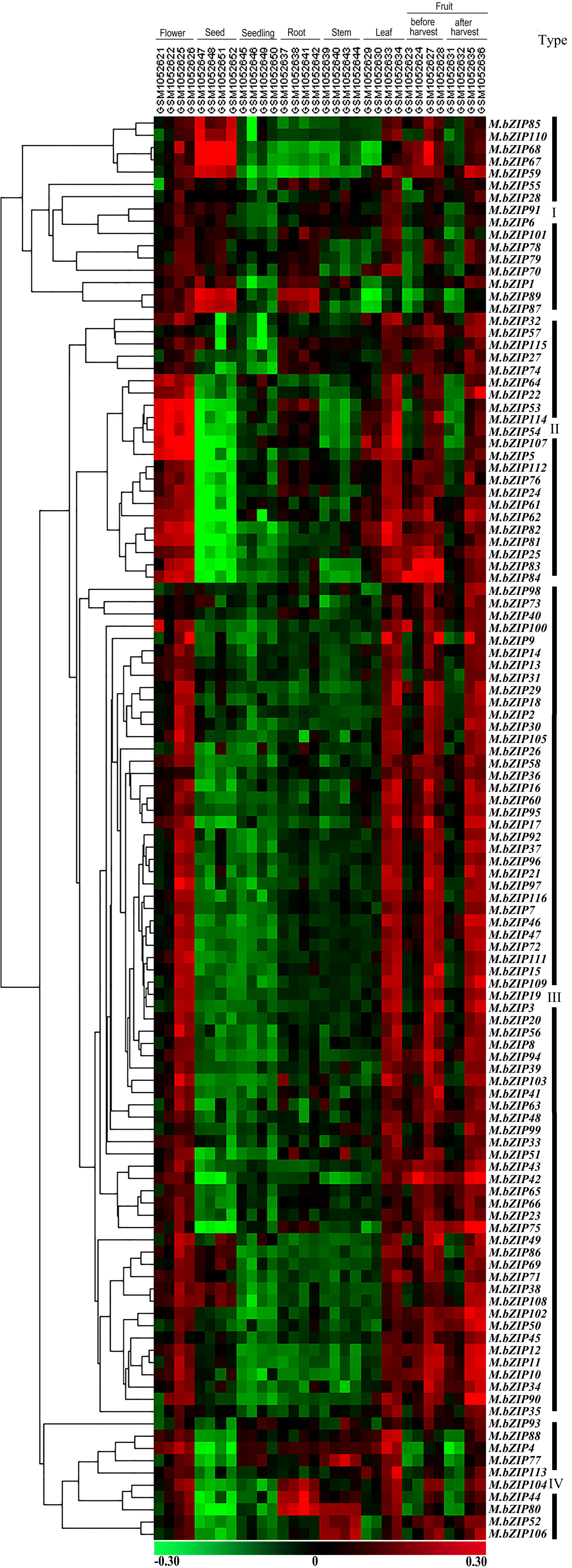
Expression patterns of *M.*bZIP genes in different organs in apple. This heatmap was generated based on the GSE expression data using cluster3.0 and treeview software. Green indicates low expression, dark indicated intermediate expression and red indicated high expression.

To verify credibility of expression profile, five *M.*bZIP genes in different types were randomly selected to check the organic expression by qRT-PCR technique using gene-specific primers (Table S6). The expression trend of *M.*bZIP genes in different organs was consistent with the expression data in the GSE42873 except *M.*bZIP52 (Figure 8). Compared with other organs, *M.*bZIP 52 has high expression in the seeds with checked by qRT-PCR, while low expression was found in expression profile in the seeds. Our results demonstrated that the expression profile of *M*.bZIP genes in different organs was reliable.

**Figure 8.**
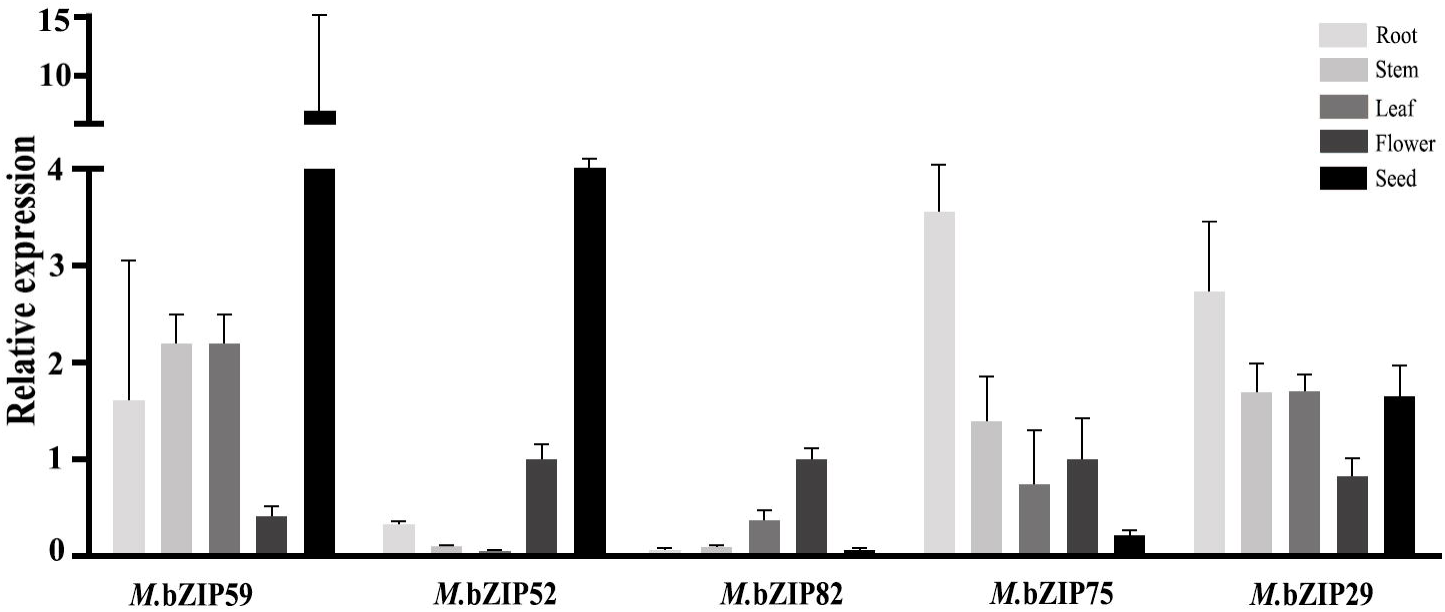
The expression changes of representive *M.*bZIP genes in flowers, seeds, roots, stems and leaves.

### 2.6. Promoter Analysis and Expression Assay of *M.*bZIP Genes under Abiotic Stress Conditions

Previous studies suggested that some bZIP proteins are involved in signaling and responses to abiotic/biotic stimuli (Kesarwani et al. 2007). To investigate the role of *M.*bZIP in stress response, the promoter sequences of fourteen randomly selected *M.*bZIP genes in different phylogenetic groups were analyzed. Total 163 *cis*-acting elements were detected, which distribution and function annotation were shown in Table S7. Important *cis*-acting elements related stress-response including ABA-response element (ABRE), dehydration-response element (DRE), low-temperature responsive element (LTRE) were widely existed in the promoters of bZIP genes as shown in Table 2. These elements have been demonstrated to bind the relative proteins to modulate stress response in other plants (Dai et al. 2007, Maestrini et al. 2009, Zhang et al. 2009, Li et al. 2012). Besides, some elements found in the promoters of *M.*bZIPs function for binding of important transcription factors, which factors were closely related to various abiotic stresses such as transcription factor MYB and MYC (Table 2). High frequence of above elements in the promoters suggested that *M.*bZIP genes were involved in the response of multiple stresses in apple.

**Table 2.** Distribution of ABRE, DRE, LTRE, MYB and MYC *cis*-acting elements in *M.*bZIP promoters.

| Group | Gene | ABRE | DRE | LRTE | MYB | MYC | Total |
| --- | --- | --- | --- | --- | --- | --- | --- |
| II | *M.bZIP16* | 1 | 0 | 1 | 24 | 21 | 47 |
| II | *M.bZIP24* | 7 | 0 | 2 | 22 | 36 | 67 |
| V | *M.bZIP29* | 4 | 0 | 1 | 14 | 18 | 37 |
| IV | *M.bZIP45* | 1 | 0 | 0 | 8 | 20 | 29 |
| VI | *M.bZIP46* | 3 | 0 | 1 | 6 | 6 | 16 |
| II | *M.bZIP49* | 6 | 0 | 0 | 14 | 14 | 34 |
| IV | *M.bZIP58* | 2 | 0 | 1 | 20 | 20 | 43 |
| III | *M.bZIP59* | 3 | 0 | 0 | 23 | 2 | 28 |
| I | *M.bZIP69* | 4 | 2 | 2 | 19 | 12 | 39 |
| II | *M.bZIP73* | 2 | 2 | 6 | 20 | 12 | 42 |
| I | *M.bZIP87* | 1 | 4 | 2 | 13 | 18 | 38 |
| I | *M.bZIP98* | 1 | 1 | 2 | 18 | 16 | 38 |
| II | *M.bZIP101* | 4 | 2 | 4 | 24 | 12 | 46 |
| V | *M.bZIP105* | 3 | 0 | 0 | 30 | 10 | 43 |

To further examine whether the expressions of *M.*bZIP gene were induced by abiotic stresses, 20% PEG and 100 μM exogenous ABA were treated to 1-year-old apple seedlings. The expression levels of seventeen genes randomly selected in different groups were checked by qRT-PCR. Most of the genes were up-regulated or down-regulated under the abiotic stress treatments (Figure 9). Among them, six *M*.bZIP genes were significantly up-regulated by ABA treatment. The relative expressions of three genes (*M*.bZIP6, *M*.bZIP45 and *M*.bZIP73) were up-regulated, approximately 3-fold (3 h), 6-fold (1 h) and 3-fold (1 h), respectively. One gene (*M*.bZIP101) exhibited decreased expression at 1 h after treatment, but then increased and reached the initial level after 3 h treatment. While the expression of ten *M.*bZIP genes showed an obvious decrease at all time points after treatment. Under osmotic stress, the expression of sixteen *M*.bZIP genes showed up-regulated and three of these (*M*.bZIP46, *M*.bZIP45 and *M*.bZIP93) were very obvious, approximately reaching 60-fold (9 h), 11-fold (3 h) and 10-fold (6 h), respectively. There was only one *M*.bZIP gene down-regulated in transcript levels at all time points after treatment with PEG. Combined with the results of the treatment of two kinds of stress, five *M*.bZIP genes increased in response to the two kinds of stress, indicated these genes play multiple roles in the regulation of the abiotic stress signaling pathways.

**Figure 9.**
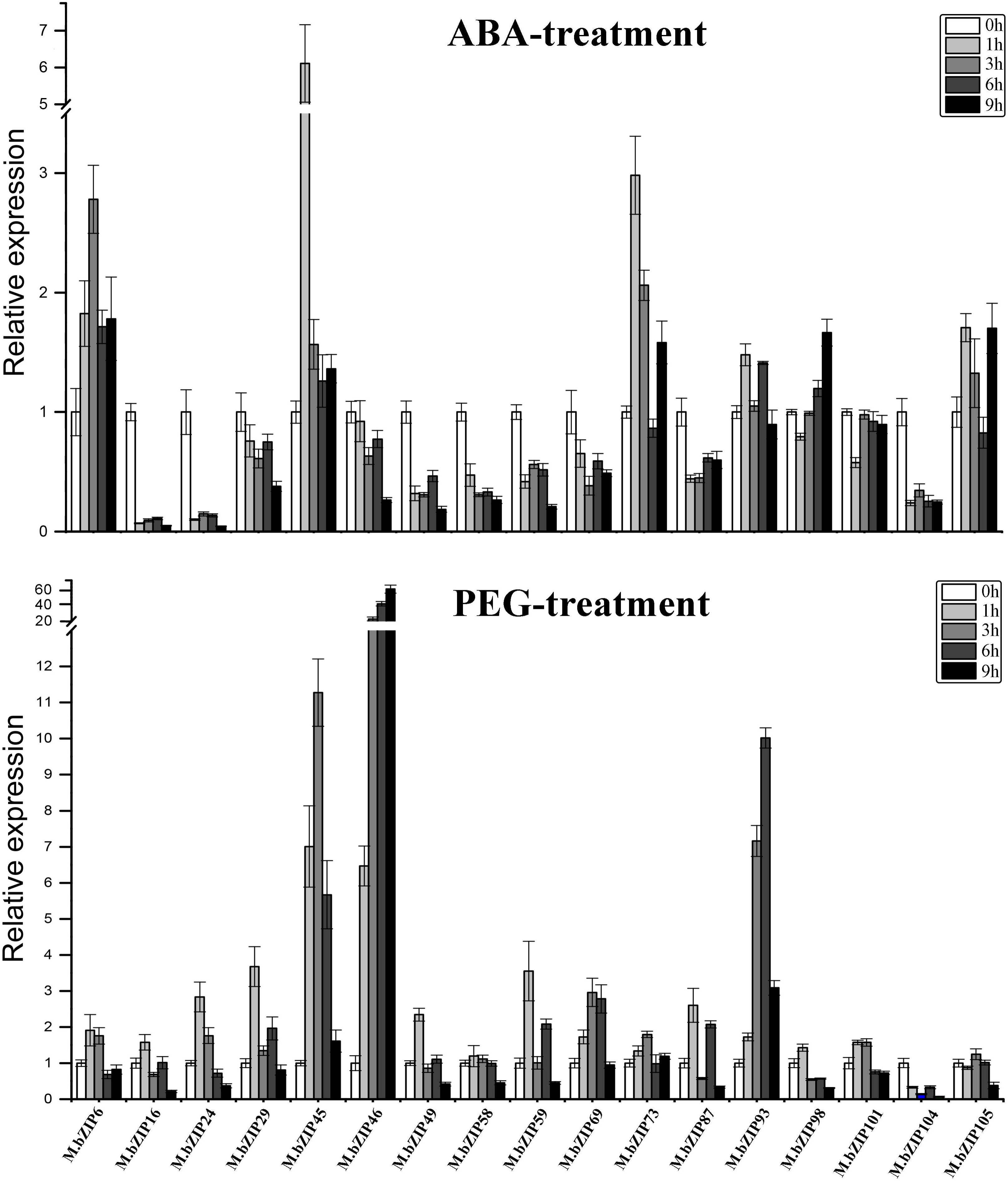
The expression changes of representive *M.*bZIP genes in different groups under ABA and PEG treatment.

## 3. Discussion

In this study, using Local Blast and Hidden Markov Model Profile, total 116 genes encoded bZIP proteins were identified in the ‘Golden Delicious’ apple genome, which was by far the largest compared with the estimates for other plants such as 76 in *Arabidopsis* (Jakoby et al. 2002) and 89 in rice (Nijhawan et al. 2008). It can be speculated that the presence of additional *M.*bZIP genes in the apple genome may reflect the great need for these genes in the complicated transcriptional regulations. Similar results were also obtained by Wang and his colleagues, when they investigated the divergence of the bZIP gene family in strawberry, peach and apple (Wang et al. 2015).

DNA-binding region of bZIPs are relatively conservative. Analyzing DNA sequence encoding this region, different splicing patterns were found. Among 116 *M.*bZIP genes, 42 members have no intron in this region, while other members with introns have four splicing patterns. In other plants, *Os*bZIPs contained seven splicing patterns and *Hv*bZIPs contained six splicing patterns (Nijhawan et al. 2008, Pourabed et al. 2015). Comprehensive considering the splicing patterns of bZIP gene family of apple, rice and barley, we find that although there are some differences of the splicing phase and the interrupted amino acids, the positions of intron are highly conserved. Splicing patterns in basic and hinge regions of bZIPs indicated that the positions of introns may be important for the evolution and divergence of the function in the *M.*bZIP family.

Compared to the gene structure, protein structure of *M.*bZIPs is directly related to the function of transcription factors. bZIP domain is the core domain of *M.*bZIP protein, however, the function diversity of the *M.*bZIP gene family may also be contributed by additional regions in bZIP proteins. In total, 25 motifs were identified in *M.*bZIPs, among them seventeen motifs appeared to contain potential phosphorylation sites (R/KxxS/T and S/TxxD/E). These phosphorylation sites were closely related to ABRE-dependent ABA signaling pathways (Fujita et al. 2013). Motif 4 and motif 7 were found to be extension of leucine zipper regions, which suggested that these two kinds of motifs may function in dimerization of *M.*bZIPs.

Gene expression patterns can provide important clues for gene function. So we also examined the expression of 116 *M.*bZIP genes in the seven organs (seeds, seedling, roots, stems, leaves, flowers and fruits) using the expression profile GSE42873. All of the *M.*bZIP genes exhibited diverse expression patterns, which represented the distinct roles of the individual *M.*bZIP genes. Interestingly, most of *M.*bZIP genes mainly expressed in leaves, flowers and fruits indicated that *M*.bZIP genes widely participated in the process of flower development and fruit ripening. We further randomly selected five *M.*bZIP genes in different types to verify the organ expression, Our results about *M.*bZIP genes in different organs by qRT-PCR were approximately consistent with the expression profile data.

Previous studies have shown that bZIP genes play pivotal roles in developmental processes and responses of multiple stresses (Izawa et al. 1994, Yin et al. 1997, Chen et al. 2012). Promoter analysis revealed that promoter regions of fourteen *M.*bZIP genes contained variety of *cis*-acting elements associated with stresses, including ABRE, DRE, LTRE, MYB and MYC. Combined with the promoter analyze, qRT-PCR was used to further investigate the expression of *M.*bZIP genes under stress. The expression revealed that some *M.*bZIP genes were sensitively responded to stress treatment. Five *M.*bZIP genes were identified to respond to two abiotic stress treatments tested, the results suggested that these genes may be stress-sensitive gene with a potential role in ABA and osmosis acclimation. Some bZIP genes, such as *M.*bZIP16 and *M.*bZIP24, exhibited opposing expression patterns under ABA and PEG treatment, which indicate that these genes are involved in the communication between different signal transduction pathways. Except *M.*bZP104, sixteen of seventeen genes which were randomly selected in different phylogenetic groups shown respond to PEG treatment. It is noteworthy that the expression of *M*.bZIP46, *M.*bZIP45 and *M.*bZIP93 showed the maximum 60-fold, 11-fold, and 10-fold increase at 9 h, 3 h and 6 h, indicating that these genes play a major role in the response to osmosis conditions. However, in the fourteen genes, the expression results from qRT-PCR did not agree perfectly with *cis*-acting elements analysis. For instance, *M*.bZIP46 contained 16 *cis*-acting elements and its relative expression level was more than 60-fold in relation to controls under PEG treatment at 9 h. whereas the value for *M*.bZIP24 carrying 67 *cis*-acting elements was only 3-fold (Table 2; Figure 9). One explanation may be that several elements lost their activities or performed in a negative way. Another explanation is that different *cis*-acting elements are different ability to initiate gene expression. Overall, our investigation about bZIP family in the apple may lead to further understanding the relationship of function and structure of bZIP family members.

## 4. Materials and methods

### 4.1. Genome-wide identification of bZIP proteins in apple

The genome and proteome sequences of *Arabidopsis* and rice were downloaded from the NCBI website. Two methods were used. For the first method, BLASTp was performed by the Bioedit program with an E value cutoff of 0.001 to search the Golden Delicious peptide database (GDR http://www.rosaceae.org/) using bZIP protein sequences in *Arabidopsis* and rice as queries. For the second method, Hmmer3.0 software was downloaded from the HMMER website (http://hmmer.janelia.org/) to perform a global search of the apple proteome. The HMM profile (PF00170) of the bZIP domain was download from the pfam website (http://pfam.xfam.orf/) (Finn et al. 2013). All of the proteins that were obtained by the two methods were submitted to the Interpro Database (http://www.ebi.ac.uk/interpro/) and smart Database (http://www.smart.embl-heidelberg.de/) to ensure the presence of bZIP domain.

### 4.2. Phylogenetic analysis of *M.*bZIP genes

116 *M.bZIP* protein sequences, 89 *Os*bZIP protein sequences and 72 *At*bZIP protein sequences were executed multiple alignments by Clustal X 1.83 program (Larkin et al. 2007). Phylogenetic trees were constructed using phylip-3.695 by the Neighbor-Joining (NJ), and the bootstrap test carried out with 1000 replications. The picture of those trees was drawn using FigTree (Verson1.4.2).

### 4.3. Chromosomal locations and gene duplications for*M.*bZIP genes

Location data were retrieved from genome annotations downloaded from the Genome Database for *Rosaceae* (GDR http://www.rosaceae.org/). The chromosome map showing the physical location of all *M.*bZIP genes was generated using revised version of MapDraw (Liu and Meng 2003). Tandem duplications were characterized as adjacent genes located within 10 predicted genes apart or within 30kbp of each other.

### 4.4. Genomic and proteome structures ofM.bZIP transcription factors

To explore the diverse exon/intron organizations of *M.*bZIP genes, we compared the predicted coding sequences of *M.*bZIP genes with their corresponding genomic sequences using GSDS software (http://gsds.cbi.pku.edu.cn) (Hu et al. 2014). The intron distribution and splicing phase within the genomic sequences were further predicted based on the alignments of the genomic sequences and cDNA full-length sequences obtained by Spidey website (http://www.ncbi.nlm.nih.gov/spidey/).

The isoelectric point (PI) of each protein was calculated by online ExPASy programs (http://www.expasy.org/tool/). The conserved motifs were identified by the online MEME analysis tool (http://meme-suite.org/). The Interpro Database was used to identify conserved functional domains in apple *M.*bZIP proteins.

### 4.5. Expression pattern of *M.bZIP* genes in apple

To study the expression pattern of *M.*bZIP genes in different organs, series matrix data from the expression profiles GSE42873 was downloaded from NCBI GEO datasets. Expression data for the identified genes were extracted from the two datasets using their unigene IDs with a Visual Basic (version6.0) script. Clustering was performed by Cluster3.0 (http://rana.IbI.gov/EisenSoftware.htm) using the hierarchical clustering method for the average linkage, a heatmap and clustering tree were constructed and viewed with Java Treeview (http://sourceoforge.net/projects/jtreeview/?source=typ_redirect).

### 4.6. Promoter analysis of *M.bZIP* genes

To investigate *cis*-acting elements in the promoter sequences of *M.bZIP* genes, 2000bp of genomic DNA sequences upstream of the transcriptional start site were obtained from the apple genome. Promoter sequences were submitted to the PLACE database (http://www.dna.affrc.go.jp/PLACE/) and cis-acting elements were obtained (Higo et al. 1999).

### 4.7. Sample preparation and total RNA extraction

Golden Delicious plants which were cultivated in the fields were selected as the experimental material for qRT-PCR analysis. The apple (Golden Delicious) seedlings were separately treated in100 μM ABA and 20% PEG6000 for 0 h, 1 h, 3 h, 6 h and 9 h. 1g leaf samples from three individual plants of stressed and controlled seedlings were harvested for total RNA extraction. The total RNA was isolated from the leaves using the CTAB procedure (Gasic et al. 2004). RNA concentrations and A_260_/A_280_ ratios were determined using a NanoDrop Spectrometer (ND-1000 Spectrophotometer, peqlab). The integrity of the RNA was detected by agarose gel electrophoresis. Qualified RNA was used for cDNA synthesis and quantitative real-time PCR.

For organ specificity expression of *M.bZIP* genes, organs of leaf, stem, root, and seed were harvested separately from one year old plants of Golden Delicious and the flowers were picked in summer. All of materials were stored at −80°C after freezing in liquid nitrogen.

### 4.8. Quantitative real-time PCR (qRT-PCR) analysis

cDNA fragments were synthesized from total RNA using the TransScrip^TM^one-step gDNA Removal and cDNA Synthesis SuperMix (TransGen Biotech, Beijing china).The gene-specific primer pairs were designed based on target gene sequences using the software Beacon Designers 8.10. The apple *actin* gene was used as an internal control. The primer sequences were listed in table S6. The qRT-PCR was carried out with a Stratagene Mx3000P thermocycler (Agilent) in a final volume of 20 µl that contained 1.4 µl cDNA, 10 µl 2*SYBR premix Ex Taq^TM^ (Takara, Shiga, Japan), 7.8 µl H_2_O and 0.8 µl (10 µM) primers. The thermal cycling conditions were as follows: 44 cycles of 95°C denaturation for 15 s, 55°C annealing for 30 s and 72°C extension for 15 s. The real-time PCR experiment was carried out at least three times under identical conditions.

## Abbreviations

ABA: abscisic acid
ABRE: abscisic acid responsive elements
bZIP gene: basic leucine zipper encoding gene
DRE: dehydration-response element
LTRE: low-temperature responsive element
qRT-PCR: quantitative real-time PCR

## Acknowledgment

This work was supported by Special Research Fund of Public Welfare of China Agricultural Ministry (201303093).

## Supplementary Material

**Table S1** *M.bZIP* genes involved in tandem duplication

**Table S2** Splicing sites and splicing phase in basic and hinge region of *M.*bZIP genes

**Table S3** Alignment sequences of bZIP domain of *M.*bZIPs

**Table S4** Conserved motifs identified from 116 *M.*bZIPs

**Table S5** Amino acid sequences alignment of the leucine zipper region of 116 *M.*bZIPs

**Table S6** List of primers used in qRT-PCR expression analysis of *M.*bZIP genes

**Table S7** C*is*-acting elements distribution in promoters of six *M.*bZIP genes and function annotation

